# Characterising the hippocampal response to perception, construction and complexity

**DOI:** 10.1101/2020.07.27.223313

**Authors:** Cornelia McCormick, Marshall A. Dalton, Peter Zeidman, Eleanor A. Maguire

## Abstract

The precise role played by the hippocampus in supporting cognitive functions such as episodic memory and future thinking is debated, but there is general agreement that it involves constructing representations comprised of numerous elements. Visual scenes have been deployed extensively in cognitive neuroscience because they are paradigmatic multi-element stimuli. However, questions remain about the specificity and nature of the hippocampal response to scenes. Here, we devised a paradigm in which we had participants search pairs of images for either colour or layout differences, thought to be associated with perceptual or spatial constructive processes respectively. Importantly, images depicted either naturalistic scenes or phase-scrambled versions of the same scenes, and were either simple or complex. Using this paradigm during functional MRI scanning, we addressed three questions: 1. Is the hippocampus recruited specifically during scene processing? 2. If the hippocampus is more active in response to scenes, does searching for colour or layout differences influence its activation? 3. Does the complexity of the scenes affect its response? We found that, compared to phase-scrambled versions of the scenes, the hippocampus was more responsive to scene stimuli. Moreover, a clear anatomical distinction was evident, with colour detection in scenes engaging the posterior hippocampus whereas layout detection in scenes recruited the anterior hippocampus. The complexity of the scenes did not influence hippocampal activity. These findings seem to align with perspectives that propose the hippocampus is especially attuned to scenes, and its involvement occurs irrespective of the cognitive process or the complexity of the scenes.

## 1. Introduction

The hippocampus makes a crucial contribution to episodic memory (Scoville & Milner, 1957), spatial navigation (O’Keefe & Dostrovsky, 1971), and a range of other cognitive domains including imagining the future (Addis, Wong, & Schacter, 2007; Hassabis, Kumaran, Vann, & Maguire, 2007), mind-wandering (Karapanagiotidis, Bernhardt, Jefferies, & Smallwood, 2016; McCormick, Rosenthal, Miller, & Maguire, 2018) and dreaming (Spano et al., 2020). Theoretical accounts differ on precisely how the hippocampus supports these diverse cognitive functions. Nevertheless, across perspectives there is a common thread, namely that its contribution involves constructing representations composed of numerous elements (Cohen & Eichenbaum, 1993; Lee et al., 2005; Yonelinas, Ranganath, Ekstrom, & Wiltgen, 2019; Hassabis & Maguire, 2007; Schacter & Addis, 2007). Visual scenes are paradigmatic multi-element stimuli and consequently have been deployed extensively to test hippocampal function in perceptual, associative, recognition, recall and imagination tasks (Aly, Ranganath, & Yonelinas, 2013; Barry, Chadwick, & Maguire, 2018; Lee, Brodersen, & Rudebeck, 2013; McCormick, Rosenthal, Miller, & Maguire, 2017). A scene is defined as a naturalistic three-dimensional spatially-coherent representation of the world typically populated by objects and viewed from an egocentric perspective (Dalton, Zeidman, McCormick, & Maguire, 2018).

Patients with hippocampal damage show scene-related perceptual, imagination and mnemonic impairments (Lee et al., 2005; Hassabis, Kumaran, Vann, & Maguire, 2007; Mullally, Intraub, & Maguire, 2012; Aly, Ranganath, & Yonelinas, 2013), and functional MRI (fMRI) studies have consistently reported hippocampal engagement during scene processing as part of a wider set of brain areas that includes ventromedial prefrontal cortex (vmPFC), parahippocampal, retrosplenial and parietal cortices (Robin, Buchsbaum, & Moscovitch, 2018; Zeidman, Mullally, & Maguire, 2015). It is unclear precisely why the hippocampus responds to scenes, and theoretical perspectives differ in the emphasis they place on specific features of scenes and their processing in order to explain hippocampal engagement, and to make inferences about its function. However, there are several gaps in our knowledge of scene processing which, if filled, may help to clarify the role of the hippocampus. Here we sought to increase our understanding by asking three questions using functional MRI (fMRI).

First, is the hippocampus especially attuned to scenes? Some accounts argue that scenes and spatial contexts merely exemplify relational processing where elements are bound together, and it is this fundamental associative processing that the hippocampus provides (Aly et al., 2013; Eichenbaum, 2006; Eichenbaum & Cohen, 2014; Erez, Cusack, Kendall, & Barense, 2016; Lee et al., 2005; Yonelinas, Ranganath, Ekstrom, & Wiltgen, 2019). A contrasting perspective proposes that the hippocampus is specifically concerned with constructing scene representations, and more so than other types of multi-feature representations (Hassabis & Maguire, 2007; Maguire & Mullally, 2013).

It is challenging to compare these differing views in a controlled way. One approach, devised by Dalton et al. (2018; see also Monk, Dalton, Barnes, & Maguire, 2020), had participants gradually build scene imagery from three successive auditorily-presented object descriptions and an imagined 3D space during fMRI. This was contrasted with constructing mental images of non-scene arrays that were composed of three objects and an imagined 2D space. The scene and array stimuli were, therefore, highly matched in terms of content and the associative and constructive processes they evoked. Moreover, the objects in each triplet were not contextually related, and for half the participants an object triplet was in a scene, and for the other half of participants it was in an array, thus controlling for semantic elements across conditions. Constructing scenes compared to arrays was associated with increased activity in the anterior medial hippocampus. Consequently, Dalton et al. (2018) concluded that it is representations that *combine* objects with specifically a 3D space that consistently engage the hippocampus, and that the anterior hippocampus may be especially attuned to constructing these scene representations (Zeidman & Maguire, 2016; Dalton & Maguire, 2017).

However, these simple scene representations are far removed from the naturalistic scenes that we experience in the real world. But if we want to examine naturalistic scenes, then it is still important to compare them to similar non-scene stimuli. One possibility, which we pursued here, is to create phase-scrambled versions of scenes (Yoonessi & Kingdom, 2008). The resulting images possess the same spatial frequency and colour scheme as the original scenes, but their phase is randomized such that any meaning is removed from the image. By combining the scenes and their phase-scrambled versions with the manipulation of a key feature of interest (complexity – more on this later), and matched cognitive task requirements, we predicted that, in line with Dalton et al. (2018), naturalistic scene stimuli would preferentially engage the anterior medial hippocampus.

If the hippocampus were more involved in scene processing, the second question we asked is whether the cognitive process engaged at the time would influence hippocampal recruitment. Lee et al. (2005; see also Lee et al., 2013; Barense, Henson, Lee, & Graham, 2010) used an odd-one-out paradigm where patients with bilateral hippocampal damage were shown three scenes from different viewpoints and were unable to select the one scene that was different from the two others. The authors interpreted this result as a scene perception deficit since all scenes were visible throughout each trial. However, it has been argued that this odd-one-out task also requires the construction of internal models of the scenes which are needed to mentally rotate the scenes in order to compare them to one another (Zeidman & Maguire, 2016). Consequently, findings such as these could reflect a hippocampal role in scene perception and/or the construction of scene imagery.

Zeidman et al. (2015) examined this issue further by having participants view visual scenes or construct scenes in their imagination, where there was the potential to be asked subsequently to hold the perceived or imagined scene in working memory. They found that perceiving scenes was associated with extensive activation in posterior hippocampus and the anterior medial hippocampus, whereas scene construction engaged the anterior medial hippocampus. This suggests that while posterior hippocampus might be particularly engaged during scene perception, anterior medial hippocampus might play a role in constructing representations of scenes, whether perceived or imagined, when there is a need to retain them in memory. While it is difficult to separate perception and construction completely, here we sought to disentangle perception and construction processes, where visual input was identical, and there was no memory requirement.

Extending previous experimental designs (Aly et al., 2013; Lee et al., 2013), on each trial participants had to examine two images that were displayed side by side and judge whether or not the two images were the same or subtly different. Images could have either a colour or layout difference. We reasoned, based on pilot testing, that a very subtle change in global colour between two images would engage participants in comparing the perceptual qualities of images to one another, while minimising the processing of layout within the image. For the constructive task, we manipulated the spatial relationships between elements within an image (see Aly et al., 2013 for a similar approach). Here, we expected that participants would focus on mentally constructing the spatial layout of one image in order to compare it to the other image. Participants were cued before each image pair whether they should focus on the colour or layout of the images. Importantly, most of the image pairs, and those that were the focus of data analysis, were identical; therefore, participants focused on colour or layout differences in the absence of visual differences. This manipulation allowed us to counterbalance the stimuli across participants so that half of the participants searched a particular image pair for colour differences and the other half of the participants searched the same pair for layout differences. We predicted that the scene colour conditions, which would most likely engage perception and would not require scene construction, would engage the posterior hippocampus. By contrast, we expected that the scene layout conditions, which were more likely to require scene construction, would recruit the anterior hippocampus.

If the hippocampus were more involved in scene processing, the third question we asked was whether its engagement would be affected by the complexity of the scenes. We used Snodgrass and Vanderwart’s (1980) definition of visual complexity as the amount of detail or intricacy in an image (see also Donderi, 2006 for a review). Thus, for example, simple scenes have a very limited number of conjunctions in the image (e.g., a deserted beach, a sky with a single bird). By contrast, complex scenes have many conjunctions (e.g., a crowded supermarket, an amusement park). Complexity is central to several perspectives on hippocampal function with high complexity, increased level of detail, number of associations or conjunctions, held to be linked to hippocampal engagement (Barense et al., 2010; Eichenbaum & Cohen, 2014; Graham, Barense, & Lee, 2010; Lee, Yeung, & Barense, 2012; Yonelinas, 2013). Therefore, the prediction of these accounts would be that complex scenes would activate the hippocampus more than simple scenes. By contrast if, as we predicted here, the hippocampus is particularly attuned to scene processing (Hassabis & Maguire, 2007; Maguire & Mullally, 2013), then any scene, simple or complex, should recruit the hippocampus.

Overall, therefore, this study had a 2×2×2 factorial design which enabled us to examine the main effects of three factors and their interactions: 1. Image type (naturalistic scenes or phase-scrambled images); 2. Task (colour or layout); and 3. Complexity (simple or complex images). To reiterate, data analysis focused on the trials where the image pairs were identical (which were the majority of trials), with stimuli counterbalanced across participants. We employed two approaches, one data driven and the other involving pre-specified contrasts. While our main focus was on the hippocampus, we examined the whole brain in order to contextualise and further inform the research questions. In addition, we recorded eye-tracking data during scanning, performed a post-scan surprise memory task to examine potential effects of incidental encoding, collected complexity ratings for the stimuli, and asked participants about the cognitive strategies they used to perform the tasks. While we had clear hypotheses, as outlined above, our paradigm also permitted evaluation of other perspectives, given the clearly contrasting predicted outcomes.

## 2. Materials and methods

### 2.1. Participants

Aligning with previous fMRI studies involving scene processing, twenty healthy, right-handed participants (8 males, mean age 27.6 years, SD 5.5, range 21-38 years) participated in the study. None had a history of neurological or psychiatric disorders. Given the pictorial nature of the stimuli, we excluded individuals engaging (as professionals, students or hobbyists) in any intensive art or design-related activities. Colour-blind individuals were also excluded as one of the tasks involved detecting subtle colour differences. All inclusion and exclusion criteria were established before data collection commenced. All participants gave informed written consent in accordance with the University College London research ethics committee.

### 2.2. Stimuli and conditions

Five hundred and six pairs of images were used in this study (26 in a pre-scan practice session, 320 during scanning, and 160 as lures in a post-scan surprise memory test). Images were all in colour, and adjusted in Adobe Photoshop CS6 to an image size of 300 dpi and cropped to the same square size (450 x 450 pixels). Given our three main factors (image type, task and complexity), there were eight main conditions: 1. Simple scene colour, 2. Complex scene colour, 3. Simple scrambled colour, 4. Complex scrambled colour, 5. Simple scene layout, 6. Complex scene layout, 7. Simple scrambled layout, 8. Complex scrambled layout. In addition, we included a number of image pairs that were of medium complexity. Their function was to act as distractors for the participants so that overall the stimuli seemed to reflect a range of complexity rather than two extremes. This resulted in four more conditions: 9. Middle scene colour, 10. Middle scrambled colour, 11. Middle scene layout, and 12. Middle scrambled layout, although the fMRI data from these middle complexity conditions were not considered as they comprised fewer stimuli than the main eight conditions. Lastly, there was also a low-level baseline task that involved viewing a fixation cross and counting from one onwards until the next cue. This was designed to allow participants to disengage from the other cognitively challenging tasks. For each of the main eight conditions, there were 25 target stimuli (that is, those with no difference between the two images in a pair) and 8 catch images (those with a difference between the two images in a pair). For each of the four middle complexity conditions, there were 10 target trials and 3 catch trials. There were 25 low-level baseline trials.

### 2.3. Image manipulations

Although we only analysed target trials in which the two images in a pair were identical and the participants identified them correctly as such, the manipulation of the catch trials was crucial to ensure that participants would engage in the different tasks, i.e., focused either on colour or layout. We, therefore, carefully created the catch trials, whereby naturalistic and scrambled as well as simple and complex images underwent the same manipulations (see Fig. 1).

**Fig 1.**
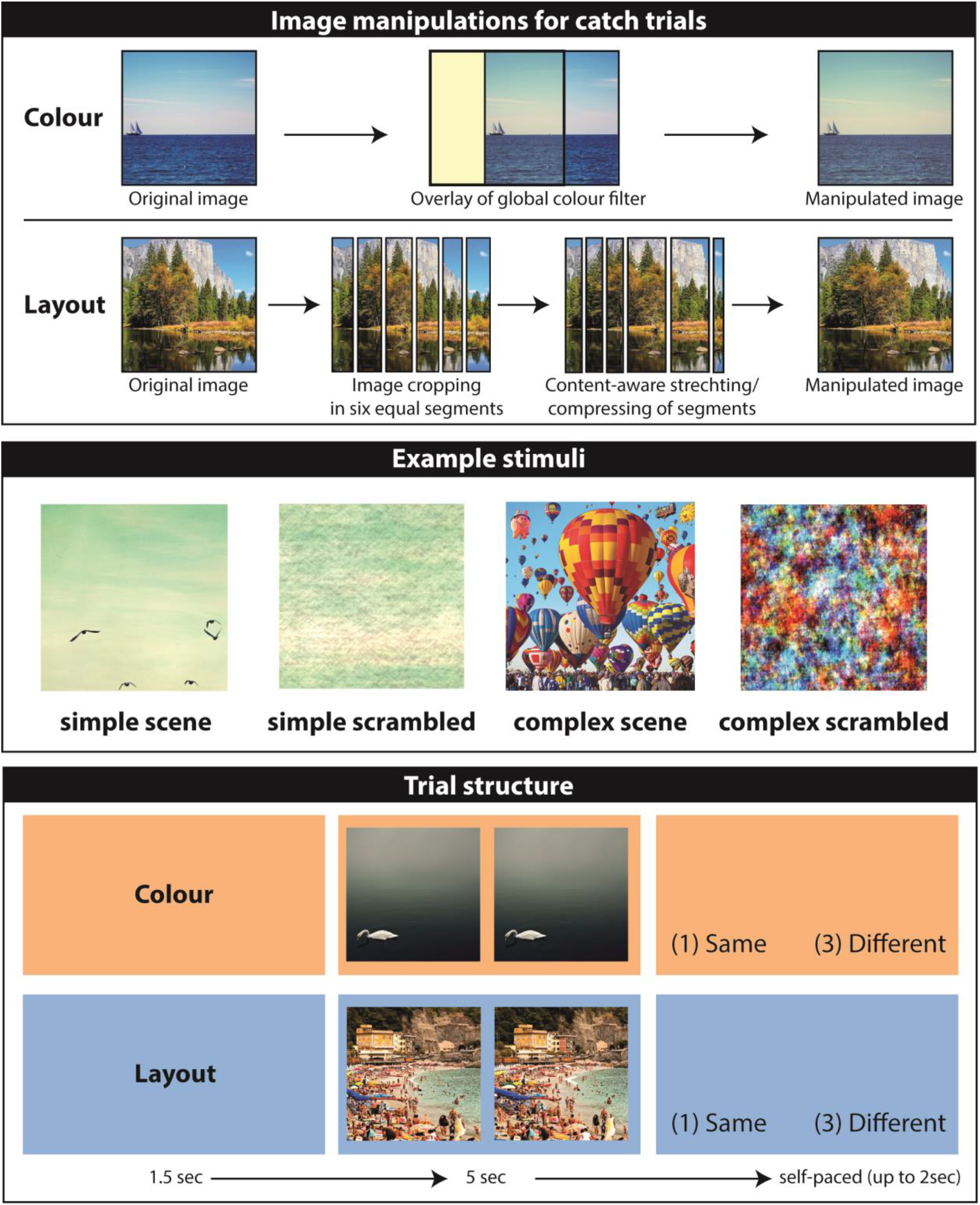
Image manipulations and experimental design. The upper panel illustrates the main image manipulations that were made in order to create the catch trials for the colour and layout conditions. The middle panel shows example scene and scrambled stimuli. Simple scenes had only a few details whereas complex scenes had many details. The scrambled versions of these scenes led to simple scrambled and complex scrambled images. The lower panels illustrate example trials, first where the participant received a cue (orange background) for 1.5sec indicating that, on this trial, they should search the upcoming image pair for a colour difference. After the cue, the image pair was displayed for 5 sec, after which there was up to 2 sec in which to make a decision. There is no difference between the images in this example. Below this is an example of a layout trial, in this case, a catch trial where there is a layout difference between the scene images.

#### 2.3.1. Colour catch trials

For the catch trials in the colour conditions, we manipulated the global colour balance of one image of a pair. For this effect, we selected one image and used Adobe Photoshop’s colour balance feature on the entire image to change the balance very slightly to either red, green or blue. Hence, when displayed, one image was shown with its original colour balance next to the altered image, creating a catch trial in which the images were different. Pilot testing (including in the MRI scanner) ensured that the colour changes were detectable but sufficiently subtle to engage the participants for the trial duration. Furthermore, the analyses reported in this study were based solely on pairs in which there were no differences between the images of a pair, yet all participants reported that they kept searching the images for colour differences for the entire trial length.

#### 2.3.2. Layout catch trials

For the catch trials in the layout conditions, we manipulated the spatial relationship between features of one image of a pair. Of note, we chose not to employ the fisheye distortion used by Aly et al. (2013) because we found that this could on occasion result in objects or lines (e.g., people, horizons), bending unnaturally. Instead, for each layout catch trial, we divided the catch image into six identical strips either vertically or horizontally. We then stretched or pinched each strip into a new dimension using Adobe Photoshop’s content-aware stretching feature which preserves lines or naturally occurring objects. Thus, whereas each strip originally occupied 1/6 of the original image, in a manipulated catch trial image, the first strip could occupy 2/6 and the sixth strip only 1/12 of the resulting image. Together, this procedure allowed us to selectively manipulate the spatial configuration of the images in a natural and global manner (i.e., the detection of errors could not be achieved by single features). At display, one image was shown with its original spatial layout next to the altered image, creating a catch trial in which the spatial layout of the images was different.

#### 2.3.3. Phase scrambling

In order to examine whether hippocampal engagement was scene-specific or not, we created phase scrambled images from the stimuli used in the scene conditions using Matlab (2014a, Mathworks), adapting a script from www.visionscience.com. This technique produced images with the same spatial frequency and colour scheme as the original scene images but because the phase was random, any meaning was removed from the images (Yoonessi and Kingdom, 2008).

#### 2.3.4. Complexity

We selected scene images that were freely available from the internet and which had varying degrees of visual complexity, basing our selection on Snodgrass and Vanderwart’s (1980) definition of visual complexity as the amount of detail or intricacy in an image (see also Donderi, 2006 for a review). There is no agreed-upon measure of visual complexity, including where to place the dividing line between simple and complex images. Much research in this domain focuses on the number of lines, objects and conjunctions that help to define the subjective feeling of image complexity (Snodgrass and Vanderwart, 1980). Here we included simple scenes that had a very limited number of conjunctions in the image (e.g., a single bird in the sky), whereas complex scenes had many conjunctions (e.g., multiple exploding fireworks in the sky). As a general rule, our complex images had over 100 conjunctions and more than 20 objects, and our simple images had under 20 conjunctions and fewer than 4 objects. Importantly, the complexity of the images did not alter significantly when simple scenes were converted into simple scrambled images and complex scenes were converted into complex scrambled images. Before commencing data collection, pilot testing endorsed our complexity classification in that simple images were judged to be visually simpler than the complex images. The participants in the fMRI study also rated scene complexity post-scanning in a very similar manner (Figure 2). Our categorisation of complexity was used in the fMRI analyses.

**Fig 2.**
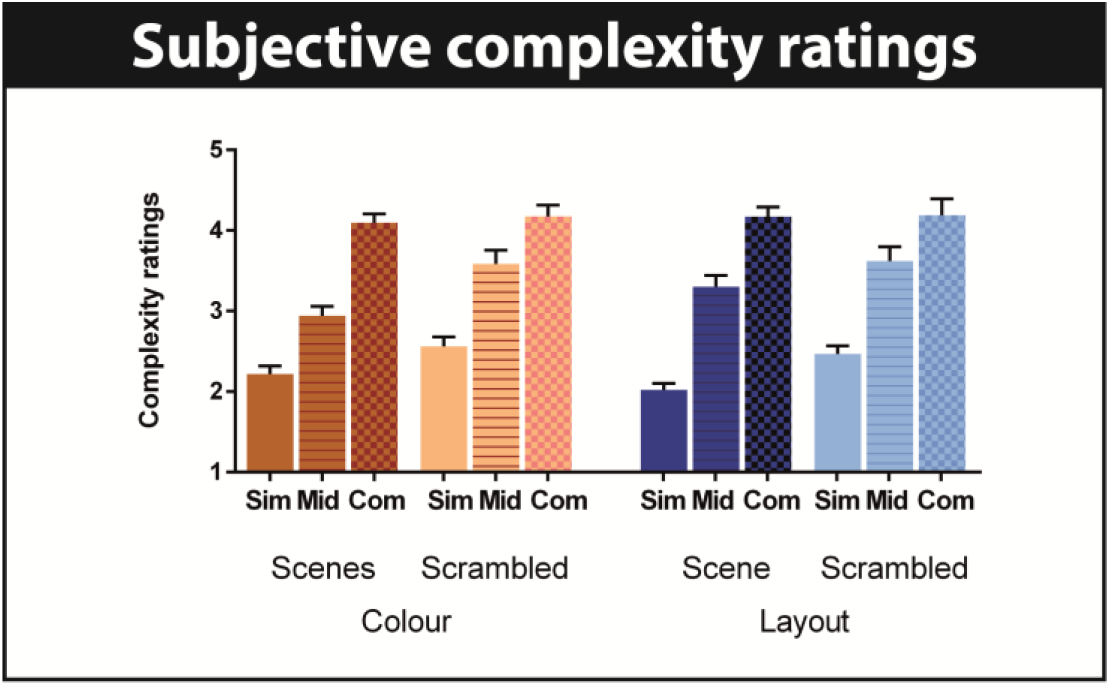
Participants’ stimulus complexity ratings. The means and standard errors of the complexity ratings made by the participants (1=very simple to 6= very complex) are shown for all conditions. Sim=simple, mid=middle, com=complex.

### 2.4. Tasks and procedure

Before scanning, participants had a short introduction and practice session. They were told that on each trial they would see a pair of images on the screen. They were instructed to look carefully at each pair because some of them would show two images that were identical whereas for others, the images would be slightly different, and that the main question they would have to answer after viewing each pair was whether the two images were the same or different. Participants were further told that a pair could differ in two ways, either in terms of the colour or the layout, and there would be cues to inform them about the type of difference they would encounter in the upcoming trial. Participants were then shown an example of a pair of scene images with one image being slightly different in colour. They were alerted to the fact that the colour change was not specific to a single object in the image but rather would affect the whole image. The participants were then shown an example of a pair of scene images with one image containing layout differences. They were told to focus specifically on the spatial relationships between features of the images and that, in the case of the example, the layout differed subtly between the two images. Participants were then shown examples of pairs with scrambled images and it was explained that in some cases, just as with the scenes, they would be asked to detect either colour or layout differences in scrambled image pairs. It was stressed to the participants that in all cases it was important to follow the cue and only look for colour differences if the cue had indicated that it was a colour trial, and only for layout differences if the cue had indicated that it was a layout trial. Moreover, they were told they should follow these instructions regardless of whether the trial involved a pair of scenes or scrambled images.

To help focus on the different tasks, pairs of images were also surrounded by an orange frame for colour trials or a blue frame for layout trials. Importantly, we counterbalanced target images (pairs of images that were identical) across participants, so that for any one image pair, 12 participants looked for colour differences and 8 for layout differences. Participants were further instructed to indicate with a button press after each pair whether they thought the pair had been the same (key 1) or different (key 3). Lastly, participants were informed that occasionally a fixation cross would appear on the screen and they were asked to empty their minds from any images and instead to count from one onwards until the next cue appeared on the screen (the low-level baseline condition).

Following these instructions, participants completed a practice session on a computer. There were 2 blocks with 13 trials each and involved stimuli that were not used in the experiment proper. The experiment was run using Cogent 2000 version 125 (Wellcome Centre for Human Neuroimaging, UCL, London, UK). Each trial started with the cue (either “Colour” or “Layout”) being displayed for 1.5 sec. Next, a pair of images was presented for 5 sec after which the decision question “Same (1) or (3) Different” appeared on the screen. Participants then responded in a self-paced manner (up to a maximum of 2 sec) by pressing the first button on the MRI button box if they thought the current pair was identical, and the third button if they thought the pair was different. After each trial of the practice session, the experimenter would give verbal feedback and if there were any mistakes, the experimenter would bring up the relevant image pair on the computer screen again after completion of the practice session for closer inspection.

After the practice session, participants were set up in the scanner, and the main experiment began. The experiment proper was completed in four blocks with 80 trials each. The trials were presented in pseudo-randomised order so that no more than two trials of the same condition were presented consecutively. The timings of the main experiment were identical to the practice session (Fig. 1). Completion of the practice and main experiment took approximately 120 minutes.

### 2.5. Eye-tracking during fMRI scanning

To examine whether and how patterns of eye movements changed depending on the image type, task or image complexity, we acquired eye-tracking data during the fMRI experiment. We used an MRI compatible Eyelink 1000 Plus (SR Research) eye-tracker during scanning and the Eyelink Data Viewer (SR Research) to examine fixation durations and fixation counts. We used the built-in online data parser of the Eyelink software whereby fixation duration was parsed automatically with fixations exceeding 100ms. The right eye was used for a 9-point grid calibration, recording and analyses. During the visual exploration of the image pairs, we recorded x and y coordinates of all fixations at a sampling rate of 1000Hz.

### 2.6. MR image acquisition

Structural and functional MRI data were acquired using a 3T Magnetom Trio scanner (Siemens Healthcare, Erlangen, Germany). The structural images were collected using a T1-weighted fast low angle shot (FLASH) sequence with 1 mm isotropic resolution (Weiskopf et al., 2013). Functional T2*-weighted images were acquired over four sessions each lasting ~15 minutes. The sequence was optimised to minimise signal dropout in the ventromedial prefrontal cortex (vmPFC) and medial temporal lobes using a slice tilt of −30 degree and a z-shim of −0.4 (Weiskopf et al., 2006). The volume TR was 3.36 sec, with a TE of 30 ms and echo spacing of 0.5 ms. Per volume, 48 slices were collected in transverse orientation, resulting in a matrix size of 64 x74 and a 3 mm isotropic voxel size. Following the first functional session, we also acquired a fieldmap with the following parameters: short TE=10 ms, long TE=12.46 ms, polarity of phase-encode blips=−1, applied Jacobian modulation=no, total EPI readout time=37 ms, in an ascending slice order.

### 2.7. Post-scan surprise memory test and complexity ratings

After the scan, participants underwent a surprise memory test. In addition, we asked participants to rate the complexity of each image. Visual complexity was explained to the participants as the level of detailedness or intricacy of an image, and an example scale of simple and complex scenes and scrambled images was provided. On a computer screen, one at a time, they saw the 320 images (scenes and scrambled) from the fMRI experiment plus 160 lures (40 simple scenes, 40 complex scenes, 40 simple scrambled, and 40 complex scrambled images). On each trial, participants responded in a self-paced manner but with a maximum of 5 sec response time to each of three questions: 1. Recognition memory: “(1) Old or (3) New”; 2. Confidence: “1=very sure, 2=somewhat sure, and 3=not at all sure”; and 3. Complexity: “(1) very simple, (2) simple, (3) middle simple, (4) middle complex, (5) complex, (6) very complex”.

### 2.8. Strategies

In a debriefing session, we asked participants to describe what strategies they had used during the experiment to search for colour or layout differences in simple and complex, scene and scrambled images. We also asked whether participants had seen any of the images before, but none had.

### 2.9. Data analysis

#### 2.9.1. Behavioural

Behavioural data collected during the fMRI scan and during the post-scan memory test were assessed using separate 2×2×2 repeated measures analysis of variance (3way-RM-ANOVA) with factor 1 being the image type with two levels (scenes, scrambled images), factor 2 being the task (colour, layout), and factor 3 being image complexity with two levels (simple, complex). Where 3way-RM-ANOVAs yielded significant effects (at p<0.05), we report the main and interaction effects. We examined significant effects further using Sidak’s multiple comparisons test and report significant results if p<0.05.

#### 2.9.2. MRI pre-processing

All MRI pre-processing was performed using SPM12 (Statistical Parametric Mapping 12; Wellcome Centre for Human Neuroimaging, London, UK). The anatomical images were segmented into grey matter, white matter and CSF maps and normalized to the Montreal Neurological Institute (MNI) template. The first five functional images were discarded to allow for signal equilibrium. Functional data were then realigned and unwarped (including distortion correction with fieldmaps) and coregistered to the anatomical image. Forward deformation fields from the anatomical image were then used to normalise the functional images into MNI space. Finally, functional images were smoothed with an 8×8×8mm kernel FWHM.

#### 2.9.3. Partial least squares (PLS) analyses

We used PLS to analyse the fMRI data. This is a multivariate, correlational technique that allowed us to examine associations between brain activity and the experimental conditions in two ways – in a contrast free, data driven manner, and also using pre-specified contrasts (Krishnan et al., 2011; McIntosh et al., 2004; McIntosh and Lobaugh, 2004). Detailed descriptions of PLS can be found elsewhere (Krishnan et al., 2011). In brief, PLS uses singular value decomposition to extract ranked latent variables (LVs) from the covariance matrix of brain activity and conditions in a data driven manner. These LVs express patterns of brain activity associated with each condition. Statistical significance of the LVs was assessed using permutation testing. In this procedure, each participant’s data was randomly reassigned (without replacement) to different experimental conditions, and a null distribution was derived from 500 permutated solutions. We considered LV as significant if p < 0.05. Furthermore, we assessed the reliability of each voxel that contributed to a specific LV’s activity pattern using a bootstrapped estimation of the standard error (bootstrap ratio, BSR). For each bootstrapped solution (100 in total), participants were sampled randomly with replacement and a new analysis was performed. In the current study, we considered clusters of 50 or more voxels with BSRs greater than 2 (approximately equal to a p < 0.05) to represent reliable patterns of activation. Of note, PLS uses two re-sampling techniques that (1) scramble the data of each participant’s conditions so that small but reliable differences between true experimental conditions can be detected, and (2) exclude whole datasets of participants, so that outliers who may drive significant effects can be detected. Therefore, even with the current sample size (n=20), we have confidence in the robustness of the results.

In a first pass, we used a mean-centred version of PLS for block fMRI data which maximises the correlation between brain data and experimental conditions in a data driven way. Importantly, this approach allowed us to examine the fMRI data without specifying a priori contrasts.

In a second pass, we used the non-rotated version of PLS for block fMRI data to specify contrasts that would test our hypotheses. While there is the potential to examine many different contrasts with this data set, we restricted our multiple comparisons to three contrast-driven PLS analyses that corresponded to our three research questions: 1. Is the hippocampus specifically engaged in scene processing? To test this, we contrasted brain activity correlated with all scene trials (regardless of task and complexity) versus all scrambled images (regardless of task and complexity). 2. If the hippocampus is specifically engaged in scene processing, does the task matter? Here, we contrasted simple and complex scene colour versus simple and complex scene layout. 3. If the hippocampus is specifically engaged in scene processing, does the complexity of the scenes matter? Here, we contrasted all simple scenes versus complex scenes (regardless of the task). To account for the multiple PLS analyses, we corrected the p-value of the LV’s using Bonferroni’s multiple comparison correction for four LV’s, resulting in a statistical threshold of p<0.017.

#### 2.9.4. Signal intensity extraction

In order to assist the reader with appreciating the specific contributions of a given brain region across all analyses and conditions, we extracted signal intensities from a number of brain regions that are typically associated with scene processing, including anterior (MNI −32 −2 −22) and posterior (−28 −36 4) hippocampus, vmPFC (2 50 −22), and occipito-temporal cortex (−6 −92 −4). Additional signal intensities (superior parietal lobule, parahippocampal gyrus and fusiform gyrus) can be found in the Supplementary Material (Fig. S1). The coordinates were chosen based on the highest boot strap ratios within these regions from the contrast driven PLS #1 (i.e., scenes versus scrambled images). Signal extraction for each condition for each participant was performed within the PLS toolbox using a sphere around the MNI coordinates of 3 adjacent voxels. Signal intensities in PLS can be positive and negative dependent on the averaged signal intensity of all the fMRI data, and do not represent percent signal change associated with experimental conditions. Therefore, these values do not reflect fMRI activation or deactivation compared to a baseline.

## 3. Results

### 3.1. In-scanner behavioural measures

In general, participants performed the in-scanner task (same or different) with high accuracy (mean of the corrected hit rate over all conditions=90.6, SD=8.8, see Table 1 for more details). The 3way-RM-ANOVA yielded a significant main effect of complexity (F(1,152)=16.5, p=0.001), as well as interaction effect of image type and complexity (F(1,152)=9.1, p=0.007), and an interaction effect of task and complexity (F(1,152)=6.5, p=0.02). However, the three way interaction between image type, task, and complexity was not significant (F(1,152)=0.2, p=0.64). Post hoc statistics revealed that these effects were driven by a lower accuracy for simple scenes than complex scenes during the colour detection task (t(152)=3.5, p=0.005). Importantly, there was no main effect of task (F(1,152)=1.4, p=0.71), nor was there an image type by task interaction (F(1,152)=3.6, p=0.07), indicating that there were no systematic differences in behavioural performance that would impact the interpretation of the fMRI data regarding our research questions. We also examined reaction times and found that there were no significant differences across conditions (all F’s(1,152)<2.6, p’s>0.11).

**Table 1:**
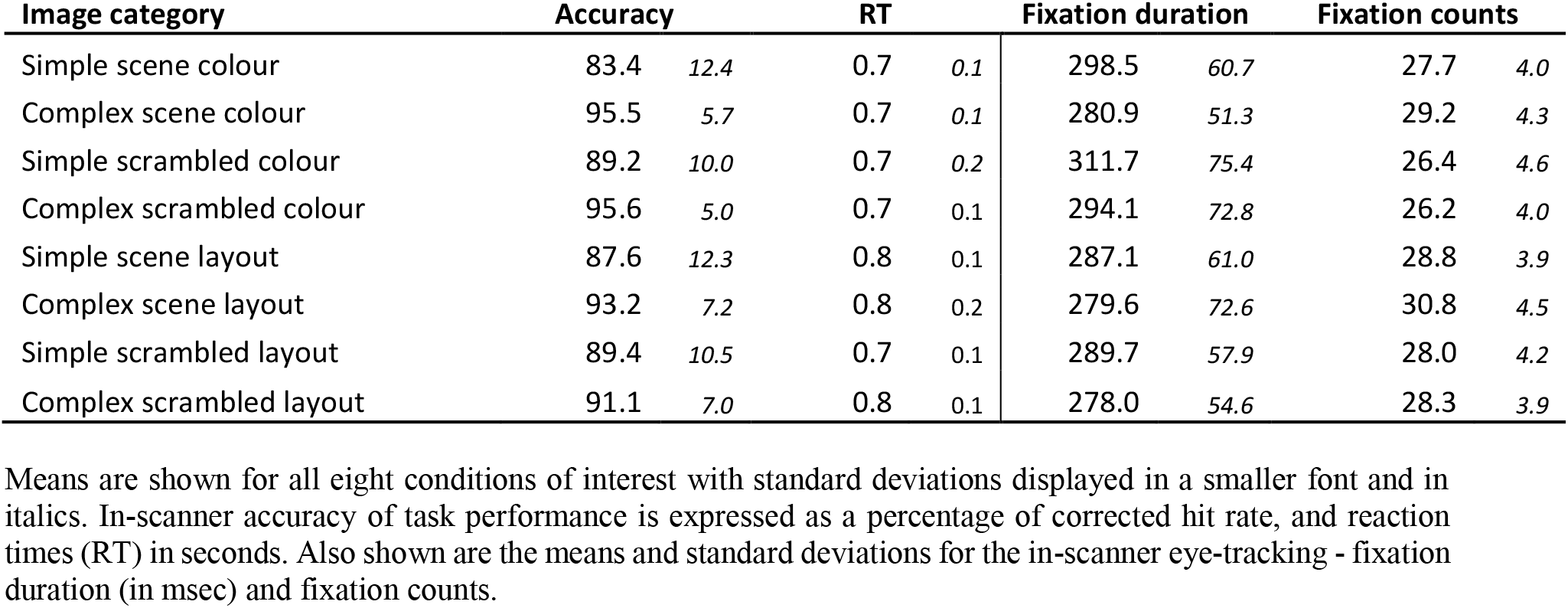
Summary of in-scanner accuracy and eye-movement measures.

### 3.2. In-scanner eye-tracking

Next, we examined eye-movements while participants were searching for colour or layout differences in simple and complex scene and scrambled images. Eye-tracking was not possible for two participants due to technical difficulties, so the following analyses are based upon data from 18 participants. We focussed on two eye-tracking measures, fixation duration and fixation counts (see Table 1 for more details).

#### 3.2.1. Fixation duration

The 3way-RM-ANOVA yielded a significant main effect of task (F(1,136)=7.1, p=0.02) and of complexity (F(1,136)=23.0, p=0.001). Post hoc analyses revealed that in general participants spent longer fixating during the colour than during the layout conditions (simple scrambled colour versus simple scrambled layout: t(17)=4.1, p<0.02; complex scene colour versus complex scene layout: t(17)=4.5, p=0.01), and on simple compared to complex images (simple scene colour versus complex scene colour: t(17)=4.0, p<0.02; simple scrambled colour versus complex scrambled colour: t(17)=4.1, p<0.02).

#### 3.2.2. Fixation counts

We observed a different pattern for fixation counts. Here, the 3way-RM-ANOVA yielded significant main effects for all three factors, image type (F(1,136)=53.5, p=0.001), task (F(1,136)=27.7, p=0.001), and complexity (F(1,136)=29.6, p=0.001). In addition, we found an interaction effect of image type and complexity (F(1,136)=16.9, p=0.001). Post hoc analyses revealed that searching images for layout differences resulted in more fixation counts than searching images for colour differences (simple scrambled colour versus simple scrambled layout: t(17)=4.2, p=0.01; complex scrambled colour versus complex scrambled layout: t(17)=4.3, p=0.002). This effect was more pronounced for scenes than scrambled images (simple scrambled colour versus simple scene colour: t(17)=4.7, p=0.003; complex scrambled colour versus complex scene colour: t(17)=9.6, p<0.0001; complex scrambled layout versus complex scene layout: t(17)=6.9, p=0.0001), and more pronounced for complex compared to simple scenes (simple scene colour versus complex scene colour: t(17)=4.0, p=0.01; simple scene layout versus complex scene layout: t(17)=4.3, p=0.007). Overall, these results indicate that participants had more fixation counts during the complex scene layout condition.

### 3.3. Post-scan surprise memory test and complexity ratings

#### 3.3.1. Memory accuracy

Due to the large number of different images and the short encoding time, recognition memory was, unsurprisingly, poor and barely exceeded chance level (mean over all conditions=60.7, SD=17.9, see Table 2 for further details). The 3way-RM-ANOVA across all conditions yielded a significant main effect of complexity (F(1, 152)=19.0, p=0.001) and an interaction effect of image type and complexity (F(1, 152)=21.4, p=0.001). Simple scrambled images for both colour (t(17)=4.7, p=0.004) and layout (t(17)=5.2, p=0.001) tasks were less well remembered than the complex scrambled images for both colour and layout conditions. This result did not come as a surprise since simple scrambled images were particularly featureless. Importantly, there was no main effect of task (F(1,152)=0.4, p=0.52), indicating that there were no systematic differences in encoding success between the colour and layout conditions that could have impacted the interpretation of the fMRI data in relation to our research questions.

**Table 2:** Summary of post-scan behavioural measures.

| Image category | Recognition memory |  | Confidence |  | Complexity |  |
| --- | --- | --- | --- | --- | --- | --- |
| Simple scene colour | 55.1 | *17.3* | 1.7 | *0.3* | 2.2 | *0.4* |
| Complex scene colour | 60.3 | *21.8* | 1.6 | *0.3* | 4.1 | *0.5* |
| Simple scrambled colour | 49.9 | *14.5* | 2.2 | *0.6* | 2.6 | *0.5* |
| Complex scrambled colour | 72.3 | *17.2* | 2.1 | *0.7* | 4.2 | *0.6* |
| Simple scene layout | 68.2 | *20.1* | 1.7 | *0.3* | 2.0 | *0.4* |
| Complex scene layout | 65.5 | *23.8* | 1.6 | *0.3* | 4.2 | *0.5* |
| Simple scrambled layout | 45.9 | *16.0* | 2.1 | *0.6* | 2.5 | *0.4* |
| Complex scrambled layout | 68.6 | *12.9* | 2.1 | *0.7* | 4.2 | *0.9* |

#### 3.3.2. Memory confidence ratings

In general, participants were not confident about whether or not they had seen a particular image in the scanner, showing that they had insight into their poor memory performance on the surprise memory test (mean over all conditions=2.0, SD=0.5, see Table 2 for further details). The 3way-RM-ANOVA across all conditions yielded a main effect of image type (F(1,152)=17.7, p<0.001), with participants, unsurprisingly, less confident about their memory judgements for scrambled images compared to scenes (simple scrambled colour versus simple scene colour: t(17)=3.2, p=0.02; complex scrambled colour versus complex scene colour: t(17)=3.4, p=0.01; complex scrambled layout versus complex scene layout: t(17)=3.6, p=0.005).

#### 3.3.3. Complexity ratings

To examine whether participants’ ratings of complexity (shown in Fig. 2) accorded with our designations, we calculated the mean complexity rating for each of our designated conditions but now based on the participants’ ratings, and entered these into a 3way-RM-ANOVA. This yielded a significant main effect of complexity (F(1,152)=428.8, p<0.0001). Post hoc analyses revealed significant differences between all simple and complex stimuli (simple scrambled colour versus complex scrambled colour: t(17)=10.6, p<0.0001; simple scene colour versus complex scene colour: t(17)=9.1, p<0.0001; simple scrambled layout versus complex scrambled layout: t(17)=12.1, p<0.0001; simple scene layout versus complex scene layout: t(17)=9.7, p<0.0001). Therefore, the participants’ ratings accorded well with our classification of complexity which was mirrored across image types and task.

### 3.4. Strategies

Also after scanning, we asked participants about how they had decided whether two images were the same or different. Generally, participants reported different strategies for colour and layout conditions, but did not report different strategies based on the image type or complexity.

For the colour task, participants indicated that they mostly focused on selected parts of the images without paying much attention to the content of the image. For example, they would compare corners, brightly lit or especially dark areas between the images. Participants indicated that they followed this strategy whether the images were simple or complex, or scenes or scrambled images. In contrast, for the layout task, participants reported examining and mentally constructing the spatial relationships within one image and then comparing these relationships to the second image. Again, participants described the same constructive strategy for simple and complex images, and for scenes and scrambled images. The strategies of each participant are summarised in the Supplementary Material (Table S1).

### 3.5. Data driven mean-centred PLS

The fMRI data driven PLS included all eight conditions of interest and revealed three significant LV’s (Fig. 3). As noted previously, this approach allowed us to examine the fMRI data without specifying a priori contrasts, enabling us to explore the dominant patterns of activity in the brain elicited by the tasks. Below we unpack each LV in turn.

**Fig 3.**
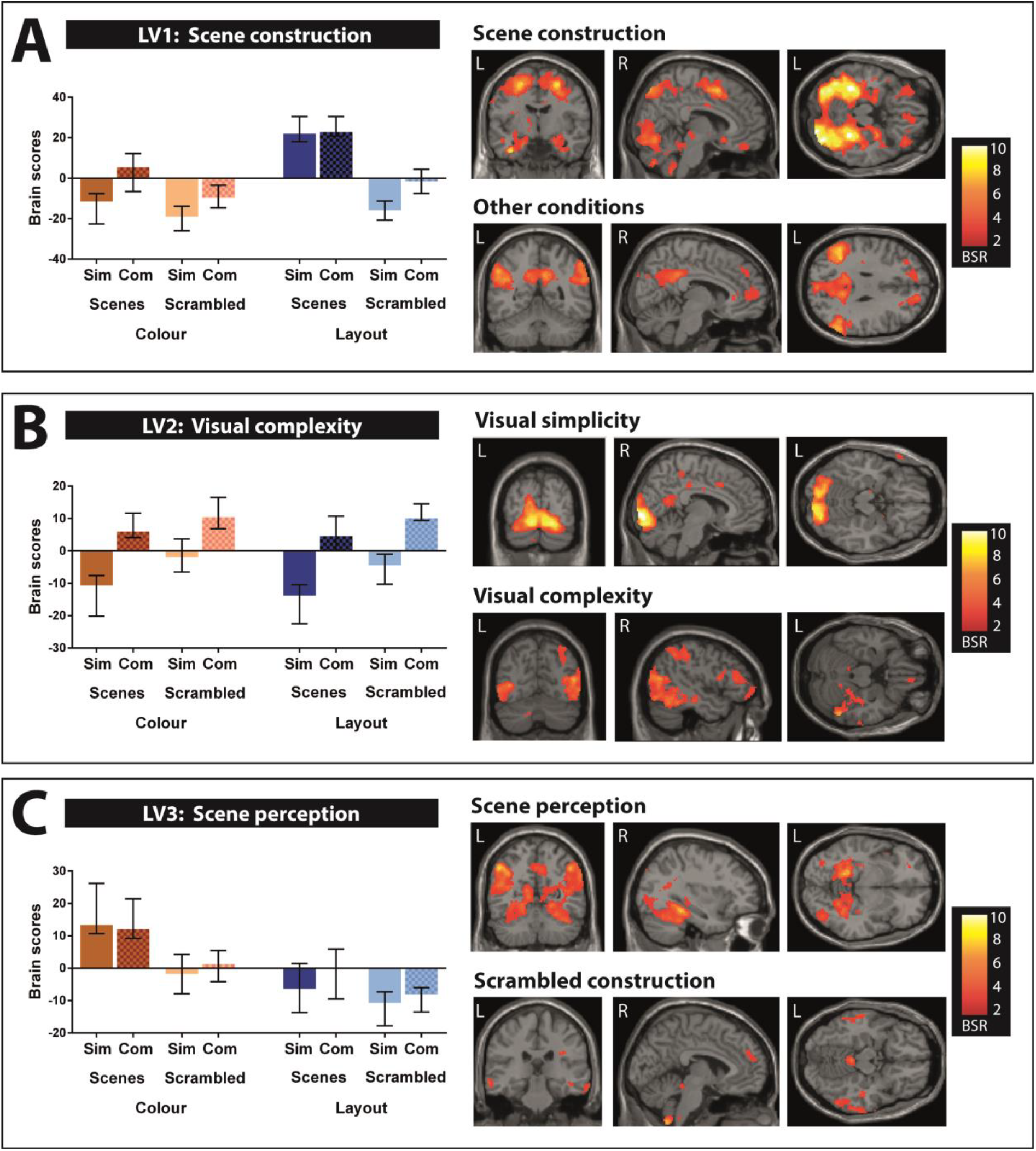
Summary of the significant latent variables (LVs) detected by the data driven PLS. Bar graphs depict means and confidence intervals for all conditions. Sim=simple, com=complex. Activations are displayed on a T1-weighted MRI scan (MNI template); L=left, R=right, BSR=boot strap ratio. (A) LV1 explained 50% of the variance. This pattern contrasted simple and complex scene layout against all other conditions, a pattern which likely reflects scene construction processes. (B) LV2 explained 15% of the variance. This pattern contrasted most simple images against complex images, regardless whether they were scenes or scrambled images, or whether participants searched images for colour or layout differences. (C) LV3 also explained 15% of the variance. This pattern highlighted simple and complex scene colour, likely reflecting scene perception processes. See also Tables S2–S4 for full details.

#### 3.5.1. LV1 – scene construction

The first significant LV (LV1, p<0.0001, explaining 50% of the variance, Fig. 3A, see Table S2 for all the brain regions that were engaged and their MNI coordinates), identified a contrast between the two conditions that we argued might be most dependent upon scene construction (i.e., simple scene layout and complex scene layout) and the other conditions. The correlated brain pattern yielded widespread activation of brain regions which are typically engaged during the construction of scene imagery, such as bilateral anterior medial hippocampus, parahippocampal gyrus, fusiform gyrus, vmPFC, bilateral precuneus and inferior parietal lobules as well as occipital cortices. Together, this finding suggests that constructing internal models of scene layouts is a dominant cognitive process associated with a distributed brain activation pattern.

#### 3.5.2. LV2 – main effect of complexity

The second significant LV (LV2, p<0.0001, explaining 15% of the variance, Fig. 3B, see Table S3 for all the brain regions that were engaged and their MNI coordinates) reflected the main effect of complexity. Interestingly, we found distinct patterns associated with simple and complex images. While simple images seemed to engage more medial posterior brain regions (e.g., medial occipital cortices), complex images engaged more lateral posterior brain regions (e.g., lateral occipital, temporal and parietal cortices). An exception to this rule was the medial subgenual vmPFC which was more activate for complex than simple images. Of note, hippocampal activity was not modulated by image complexity. Overall, this LV suggests that complexity as a general image feature engages mostly posterior visual brain regions. Nevertheless, one has to interpret this result with caution since this main effect reflects a combination of multiple, very different conditions. In our follow-up contrast driven PLS analyses, we specified more tailored contrasts to examine the effect of stimulus-specific complexity.

#### 3.5.3. LV3 – scene perception

The third significant LV (LV3, p<0.02, explaining 15% of the variance, Fig. 3C, see Table S4 for the relevant brain regions and their MNI coordinates) identified a contrast between conditions that we argued mostly depend upon scene perception (i.e., simple scene colour and complex scene colour) and the search for layout differences in scrambled images (i.e., simple scrambled layout and complex scrambled layout). While searching scrambled images for layout differences involved mainly lateral temporal, parietal and dorsomedial prefrontal cortices, the associated brain pattern for examining scenes for colour differences involved the posterior hippocampi, as well as several brain regions along the ventral visual pathway, such as the fusiform gyrus, parahippocampal gyrus and the inferior temporal gyrus. Of note, a number of regions previously found to be associated with scene construction, such as the anterior medial hippocampi and the vmPFC, were absent for this brain activity pattern. Again, this LV resulted from a data driven method, hence the interpretation of contrasting conditions involving the search for colour differences in scenes against scrambled layout is not immediately clear. Therefore, in a second pass, we conducted non-rotated PLS analyses where conditions that involved searching scenes for colour or layout differences were directly contrasted.

### 3.6. Contrast driven non-rotated PLS

We had three specific research questions which we focussed on in the analyses described below: 1. Is the hippocampus specifically engaged in scene processing in this experiment?; 2. If hippocampal engagement is specific to scene processing, does the task (i.e., colour or layout) matter?; and 3. If the hippocampus is responsive during scene processing, does the complexity of the scenes matter?

#### 3.6.1. Contrast driven PLS #1: Is the hippocampus specifically engaged in scene processing?

To answer this question, we contrasted all four conditions involving scene processing (simple scene colour, complex scene colour, simple scene layout, and complex scene layout) against the other four conditions of scrambled images (simple scrambled colour, complex scrambled colour, simple scrambled layout, and complex scrambled layout). The PLS revealed a significant LV (p<0.0001, Fig. 4A, see Table S5 for the relevant brain regions and their MNI coordinates) separating all scene conditions from all scrambled conditions. Confirming our hypotheses, the brain pattern associated with scene processing included bilateral hippocampi (both anterior and posterior segments), as well as the usual ventral visual brain regions that are typically associated with scene processing. In addition, the subgenual vmPFC also showed greater activation during scene processing than for the scrambled conditions. In contrast, processing of scrambled images was associated with a much more restricted pattern of brain activity that included engagement of the precuneus and the anterior cingulate cortex.

**Fig 4.**
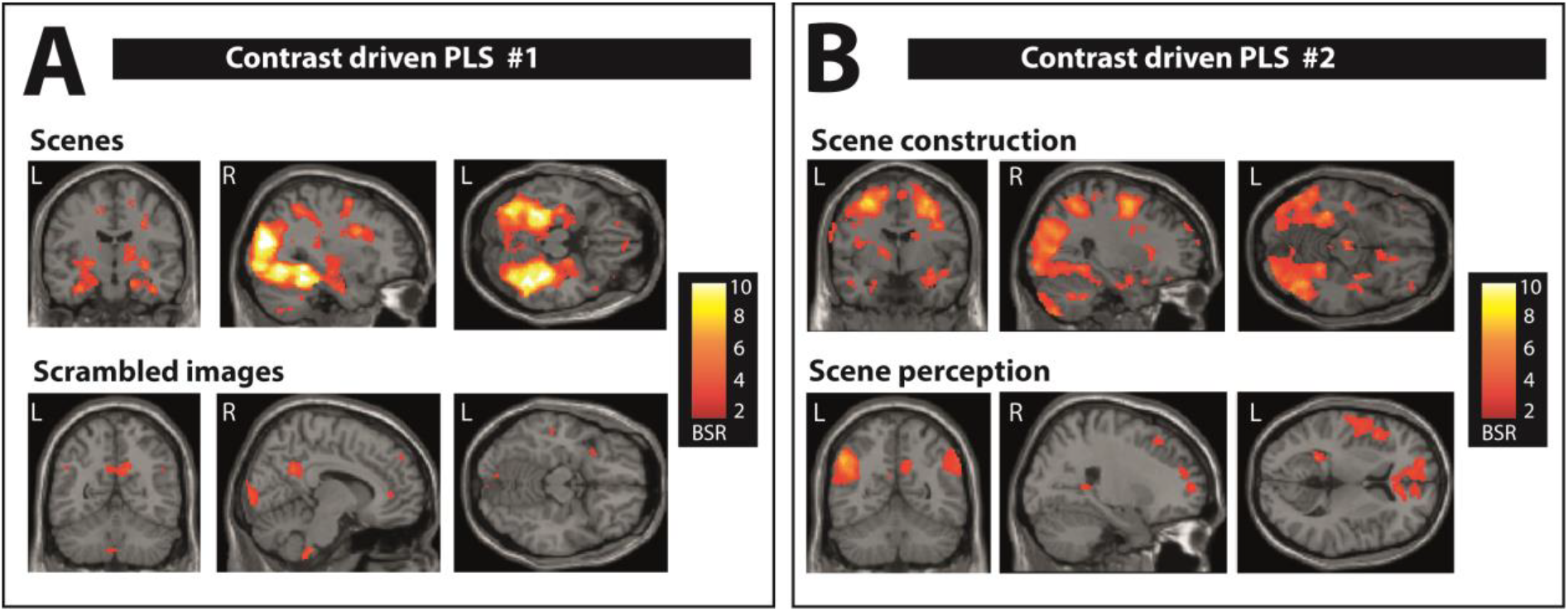
Brain activity patterns associated with the contrast driven PLS. Activations are displayed on a T1-weighted MRI scan (MNI template); L=left, R=right, BSR=boot strap ratio. (A) The brain activity pattern associated with simple and complex scene colour and layout conditions, shown in the upper panel, reflected the well-established set of brain regions associated with scene processing including increased activity in the vmPFC, bilateral hippocampus and along the ventral visual stream. This is in contrast to the brain pattern associated with simple and complex scrambled colour and layout conditions, shown in the lower panel, which included lateral and medial parietal cortices and anterior cingulate cortex. (B) The brain activity pattern associated with simple and complex scene layout (most likely depending on scene construction), shown in the upper panel, included increased activity along the ventral visual stream, bilateral anterior medial hippocampus and vmPFC. The brain pattern associated with simple and complex scene colour (most likely depending on scene perception), shown in the lower panel, included medial and lateral parietal cortices and anterior cingulate cortex. In addition, we observed increased bilateral posterior hippocampal activity for the scene colour conditions. See also Tables S5–S6 for full details.

**Fig 5.**
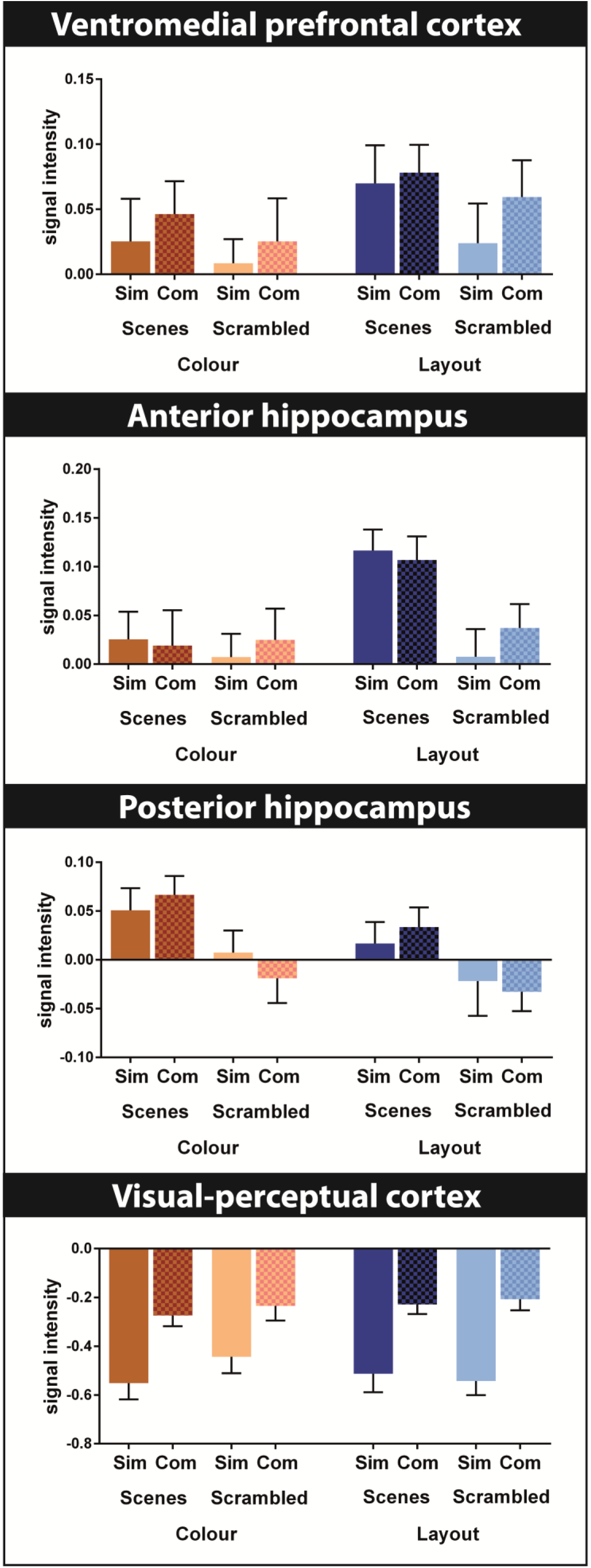
Extracted signal intensities from brain regions associated with scene processing. Bar graphs depict means and standard errors of the eight conditions for the vmPFC, anterior and posterior hippocampus and occipito-temporal cortex. Sim=simple, com=complex. These regions were chosen based on the highest bootstrap ratio in the associated contrast driven PLS #1 analysis. Of note, signal intensities are compared to an arbitrary fMRI baseline, hence negative values do not necessarily represent deactivations. Activity in vmPFC had a multifaceted pattern, reflecting a preference for scenes, especially scene layout conditions, whilst also keeping track of the visual complexity of all stimuli. The anterior and posterior hippocampus had distinct patterns of activity that were more clear-cut, with the former engaged by conditions associated with scene construction, and the latter by conditions associated with scene perception. Activity within the visual cortex reflected predominantly visual complexity. Additional signal intensities (superior parietal lobule, parahippocampal gyrus and fusiform gyrus) are provided in the Supplementary Material (Fig. S1).

#### 3.6.2. Contrast driven PLS #2: If the hippocampus is specifically engaged in scene processing, does the task matter?

Given that in the first contrast we identified greater hippocampal involvement in scene processing compared to scrambled images, we next asked whether the brain activity patterns, including hippocampal engagement, was specific to conditions that most likely involve scene construction. We, therefore, contrasted conditions that involved searching scenes for layout differences (simple scene layout and complex scene layout) against those involving the search for colour differences in scenes (simple scene colour and complex scene colour), which most likely involved scene perception. The resulting LV was highly significant (p<0.0001, all conditions contributing, Fig. 4B, see Table S6 for all relevant brain regions and their MNI coordinates). The hippocampus was among the areas engaged during both scene construction and scene perception. Interestingly, there was a clear dissociation between its anterior and posterior segments in terms of their responsiveness to the different types of scene processing. Whereas searching for colour differences engaged the posterior hippocampus, searching scenes for layout differences engaged bilateral anterior hippocampus. Furthermore, while the scene colour conditions engaged the precuneus and angular gyrus, as well as anterior cingulate cortex, the scene layout conditions were associated with several brain regions along the ventral visual pathway and superior parietal lobule. In addition, the subgenual vmPFC was also more activated during the scene layout compared to scene colour conditions.

#### 3.6.3. Contrast driven PLS #3: If the hippocampus is specifically engaged in scene processing, does the complexity of the scenes matter?

Here, we contrasted simple (simple scene colour and simple scene layout) versus complex scenes (complex scene colour and complex scene layout). However, this PLS analysis did not reveal a significant LV (p=0.06 with the Bonferroni cut-off being p<0.017).

#### 3.6.4. Regional signal intensities

We extracted signal intensities from several of the brain which had the highest bootstrap ratios in the contrast driven PLS #1 analyses. The resulting graphs merely illustrate what has already been detected by the PLS pattern and do not offer any new information per se, however, they do provide a convenient overview of the activity patterns across all eight experimental conditions. Activity in vmPFC (x y z: 2 50 −22) had a multifaceted pattern, reflecting a preference for scenes, especially scene layout conditions, whilst also keeping track of the visual complexity of all stimuli. The anterior (−32 −2 −22) and posterior (−28 −36 4) hippocampus had distinct patterns of activity that were more clear-cut, with the former engaged by conditions tapping into scene construction, and the latter by conditions tapping into scene perception. Activity within the occipito-temporal cortex (−6 −92 −4) reflected predominantly visual complexity. Additional signal intensities (superior parietal lobule, parahippocampal gyrus and fusiform gyrus) can be found in the Supplementary Material (Fig. S1).

## 4. Discussion

In this study we sought to deepen our understanding of hippocampal processing by addressing three questions: 1. Is the hippocampus recruited specifically during scene processing? 2. If the hippocampus is active in response to scenes, does the task, and the cognitive process it likely engages, influence its activation? 3. If the hippocampus is upregulated during scene processing, does the complexity of the scenes affect its response? We found that, compared to phase-scrambled versions of the scenes, the hippocampus was more responsive to scene stimuli. Moreover, there was a clear distinction in terms of which parts of the hippocampus were engaged, with conditions that likely relate to scene perception associated with the posterior hippocampus and conditions that tend to depend on scene construction involving the anterior hippocampus. The complexity of the scenes did not influence hippocampal activity. We discuss each of these results in turn.

### 4.1. The hippocampus is upregulated during scene processing

The hippocampi (anterior and posterior segments) were more activated for scenes than scrambled images. This echoes previous work using simplified representations of scenes and non-scene arrays which also showed preferential engagement of the hippocampus for scenes (Dalton et al., 2018; Monk, Dalton, Banes, & Maguire, 2020), in that case also controlling for semantic elements. Our findings align in particular with accounts of hippocampal function that propose the hippocampus is especially attuned to scene processing (Hassabis & Maguire, 2007; Maguire & Mullally, 2013). We acknowledge that it is challenging to devise non-scene stimuli for comparison with naturalistic scenes. Here, we contrasted the scenes with images that were phase-scrambled versions of the same scenes, thus preserving the spatial frequency and colours. The scenes and scrambled stimuli were rated comparably by participants in terms of the various levels of complexity. There was no main effect of image type for the same/different judgements during scanning, or in eye movement fixation duration. The pattern of memory performance in the surprise post-scan test was similar, in particular for complex scenes and complex scrambled stimuli. Moreover, the strategies participants reported using during the tasks did not differ as a function of stimulus type. Nevertheless, despite all of these similarities, the hippocampus was engaged preferentially for scenes.

It is perhaps not surprising that scenes are particularly stimulating for the hippocampus, as they mirror how people experience and perceive the world. In addition, scenes are a highly efficient means of packaging information. Clark et al. (2019) recently reported that the ability to construct scene imagery explained the relationships between episodic memory, imagining the future and spatial navigation task performance. The prominence of scenes was further emphasised in another study involving the same sample, where the explicit strategies people used to perform episodic memory recall, future thinking and spatial navigation tasks was assessed (Clark, Monk, & Maguire, 2020). In each case, the use of scene visual imagery strategies was significantly higher than for all other types of strategies (see also Andrews-Hanna, Reidler, Sepulcre, Poulin, & Buckner, 2010). The apparent utility and prevalence of scene processing has led to the suggestion that scene imagery may be the currency of cognition (Maguire & Mullally, 2013).

### 4.2 A hippocampal distinction between scene perception and scene construction

Given that scenes activated the hippocampus more than scrambled versions of the same scenes, we next considered whether the task, and by extension the cognitive process that was likely being engaged, influenced hippocampal recruitment. As outlined previously, the conditions that involved searching images for layout differences were held to tap into constructive cognitive processes, while examining images for colour differences involved perceptual processes. The scene construction and perception tasks were well-matched in a number of respects. Importantly, we counterbalanced images across participants so that all analyses dissociating perception and construction processes were conducted on the same images. The accuracy during scanning where participants correctly identified pairs of images as being identical did not differ between colour and layout trials. In addition, recognition memory assessed during a surprise memory task after scanning was similar for both tasks, indicating that there was no disparity in terms of incidental encoding. Despite these similarities, there were differences in the hippocampal response to perception and construction.

A striking result of the data driven mean centred PLS (LV1) was a clear anterior medial hippocampus preference for the two conditions thought to weigh most heavily on scene construction, namely the tasks involving the processing of simple and complex scene layouts. This finding accords with the numerous previous reports associating scene construction with the anterior hippocampus (reviewed in Zeidman & Maguire, 2016; see also Zeidman et al., 2015a,b; Dalton et al., 2018; Dalton & Maguire, 2017). By contrast, another data driven LV, this time involving the posterior hippocampus, was associated with conditions relating to the perception of scenes, simple and complex, with no evidence of the anterior hippocampus in this brain pattern. A direct contrast between the scene layout and scene colour conditions confirmed that the two processes involved different hippocampal segments, anterior and posterior respectively. These findings are in line with research stressing functional dissociations along the anterior-posterior axis of the hippocampus during scene construction and scene perception (e.g., Zeidman et al., 2015a; Zeidman and Maguire, 2016), and more generally (Poppenk, Evensmoen, Moscovitch, & Nadel, 2013; Sekeres, Winocur, & Moscovitch, 2018; Strange, Witter, Lein, & Moser, 2014).

The medial hippocampus is well suited to scene-based cognition given its anatomical (Dalton & Maguire, 2017) and functional (Dalton et al., 2019a,b) connectivity with regions in the parieto-medial temporal pathway. While the resolution of the current fMRI data do not permit hippocampal subfield analyses, it is important to acknowledge that subfields might further differentiate between perceptual and constructive processes, as well as scenes and other types of stimuli. For example, recent high-resolution fMRI studies indicate that the anterior medial portion of the subiculum or pre-/parasubiculum may be specifically engaged during scene construction (Zeidman et al., 2015b; Hodgetts et al., 2017; Dalton et al., 2018). In contrast, several previous fMRI studies showed that posterior hippocampal activity was associated with tasks involving perceptual discrimination between visually similar scenes (Aly et al., 2013; Barense et al., 2010; Lee, Buckley, et al., 2005; Lee, Bussey, et al., 2005). The posterior hippocampus has strong anatomical and functional connections with the ventral visual stream and early visual cortices (Chadwick, Mullally, & Maguire, 2013; Kahn, Andrews-Hanna, Vincent, Snyder, & Buckner, 2008). Therefore, it may be that the posterior hippocampus is involved in guiding ongoing scene perception while the anterior hippocampus supports online construction into a coherent mental model of the world.

Interestingly, eye-tracking data recorded during scanning revealed a difference between the colour and layout tasks. Whereas the colour conditions were associated with longer fixation durations, the layout conditions had more fixation counts. This effect was most pronounced during processing of scenes, especially complex scenes. These results generally align with extant studies linking rapid visual sampling to the construction of mental events (El Haj & Lenoble, 2017; Hannula & Ranganath, 2009; Liu, Shen, Olsen, & Ryan, 2017). Of note, since there were no recognition memory differences between colour and layout trials, our eye-tracking results speak against a proposal that exploratory visual sampling is purely memory-guided (Voss, Bridge, Cohen, & Walker, 2017). Rather, our findings seem to indicate that the pattern of eye-movements relates to an interaction between the dominant cognitive process and the image type (naturalistic versus scrambled, complex versus simple) during a particular task.

We also asked participants directly about their cognitive strategies during the colour and layout tasks. All participants reported distinct cognitive approaches to searching image pairs for colour versus layout differences. They indicated that they mostly focused on selected parts of the images without paying much attention to the content of the image during colour trials. For example, they would compare corners, brightly lit or especially dark areas between the images. In contrast, for the layout task, participants reported examining and mentally constructing the spatial relationships within one image and then comparing these relationships to the second image, also without paying much attention to the content of the scenes. As already noted, the images in the scene (and scrambled) conditions were counterbalanced across participants, such that half the participants searched a specific scene for colour differences and the other half of the participants searched the same scene for layout differences. Hence, the same scene content (and semantic meaning) was present in both conditions, and so is unlikely to explain our results. Furthermore, we have recently shown that while patients with hippocampal damage have difficulty detecting spatial-constructive impossibilities in scenes (e.g., an endless stair case), they did not have any problem detecting semantic impossibilities in scenes (e.g., an elephant with butterfly ears, see McCormick et al., 2017).

Overall, therefore, our results suggest that both cognitive processes, scene perception and scene construction, engaged the hippocampus, but with long-axis differences in the portion most involved. The next question we addressed was whether or not the complexity of scenes affected hippocampal recruitment.

### 4.3. No effect of scene complexity on hippocampal engagement

In the current study, we operationalised visual complexity in terms of the amount of detail or intricacy of an image (Snodgrass & Vanderwart, 1980; see also Donderi, 2006), and the participants showed high agreement with our designations of simple, middle and complex images. A number of current hippocampal theories argue that visual complexity (or the number of associations or conjunctions), is an important driver of hippocampal activity (Eichenbaum & Cohen, 2014; Lee et al., 2012; Yonelinas, 2013). Thus, more complex images should evoke greater hippocampal response compared to simpler images. This is in contrast to another perspective that suggests a primary function of the hippocampus is to construct scene imagery, irrespective of whether the scenes are simple or complex (Hassabis & Maguire, 2007; Maguire & Mullally, 2013). Complexity might also be relevant for another issue, namely the assertion by Poppenk, Evensmoen, Moscovitch, & Nadel (2013; see also Brunec et al., 2018) that representations in the hippocampus vary from fine to coarse grained in a posterior to anterior direction. This leads to the prediction that complex scenes should engage the posterior hippocampus.

However, the data driven and contrast driven analyses showed that hippocampal activity was not influenced by scene complexity. Moreover, the cognitive strategies used by participants did not differ for simple and complex scenes. Instead, complexity as a general image feature engaged mostly posterior visual brain regions. The simple and complex scenes in this study differed vastly in terms of their complexity, and so we believe they offered a credible test of the effect of complexity in terms of naturalistic scenes. Our finding accords with the view that the hippocampus processes scenes regardless of whether they are simple or complex (Hassabis & Maguire, 2007; Maguire & Mullally, 2013). How perspectives advocating complexity as a key feature of hippocampal processing (e.g., Lee et al., 2012; Yonelinas, 2013), or the view that specifically fine grained (e.g., complex) representations would engage posterior hippocampus (Poppenk, Evensmoen, Moscovitch, & Nadel, 2013; Brunec et al., 2018), which they did not, can account for our results is unclear. It may be that such theories need to define notions of complexity more precisely, to stipulate the specific processes or features of real-world perception and mental representations that might be subject to this purported effect. Certainly we can conclude from the current study that the number of objects, associations and conjunctions in naturalistic scenes did not influence hippocampal engagement.

### 4.4. Beyond the hippocampus

The focus of the current study was the hippocampus. Nevertheless, our analyses revealed that the hippocampus was part of a wider set of activated brain areas, including many that have been previously implicated in scene and event processing, such as the parahippocampal and fusiform gyri, and parietal cortex. Among these areas the vmPFC had perhaps the most interesting profile. While it was recruited to a greater extent during layout compared to colour conditions, it also seemed to be more responsive to scenes than scrambled images and to complex than simple stimuli. We speculate that this result might suggest that the vmPFC is a hierarchically superordinate structure that keeps track of scene processing (for a related idea see Robin & Moscovitch, 2017; Sekeres et al., 2018). In fact, we have suggested that the vmPFC may initiate scene construction processes in the hippocampus (Ciaramelli, De Luca, Monk, McCormick, & Maguire, 2019; McCormick, Ciaramelli, De Luca, & Maguire, 2018; see also De Luca, McCormick, Mullally, Intraub, Maguire, & Ciaramelli, 2018; De Luca, McCormick, Ciaramelli, & Maguire, 2019). In support of this proposal, recent magnetoencephalography studies have found vmPFC activity preceded and then drove that of the hippocampus during both scene imagination (Barry, Barnes, Clark, & Maguire, 2019; Monk, Dalton, Barnes, & Maguire, 2020) and the recall of autobiographical memories (McCormick et al., 2020).

### 4.5. Conclusions

In this study we sought to probe hippocampal function by manipulating three factors. We found evidence that the hippocampus was engaged by naturalistic scenes compared to scrambled images. Furthermore the posterior hippocampus was activated to a greater extent during tasks relating to scene perception and the anterior hippocampus during tasks associated with scene construction, regardless of the complexity of scenes. In-scanner task performance and incidental encoding could not explain these findings. Overall, these results seem to fit best with the view that the hippocampus may be attuned to processing scenes, be they simple or complex (Hassabis & Maguire, 2007; Maguire & Mullally, 2013). This conclusion could be investigated further in future studies by testing patients with bilateral hippocampal damage, whereby the prediction would be that they should be impaired on tasks involving scenes, be they simple or complex, but unimpaired on tasks involving scrambled stimuli.

## Acknowledgements

We are grateful to David Bradbury and Eric Featherstone for technical assistance.

## Supplementary Material

Supplementary data are available for this article.

## CRediT authorship contribution statement

**Cornelia McCormick:** Conceptualization, Methodology, Investigation, Formal analysis, Writing - original draft, Writing - review & editing. **Marshall A. Dalton:** Conceptualization, Writing - review & editing. **Peter Zeidman:** Conceptualization, Writing - review & editing. **Eleanor A. Maguire:** Conceptualization, Methodology, Funding acquisition, Supervision, Formal analysis, Writing - original draft, Writing - review & editing.

## Notes

No part of the study procedures or analyses was preregistered prior to the research being conducted. We report how we determined our sample size, all data exclusions (if any), all inclusion/exclusion criteria, whether inclusion/ exclusion criteria were established prior to data analysis, all manipulations, and all measures in the study. The raw neuroimaging data cannot be shared publicly due to ethical restrictions relating to General Data Protection Regulation. Data will be released to researchers on the following conditions: approval from the local research ethics committee and with appropriate safeguards to protect from identification of individuals. The stimuli and analysis software are already in the public domain. Any questions can be sent to the corresponding author via email.

## Supplementary Material

**Figure S1.**
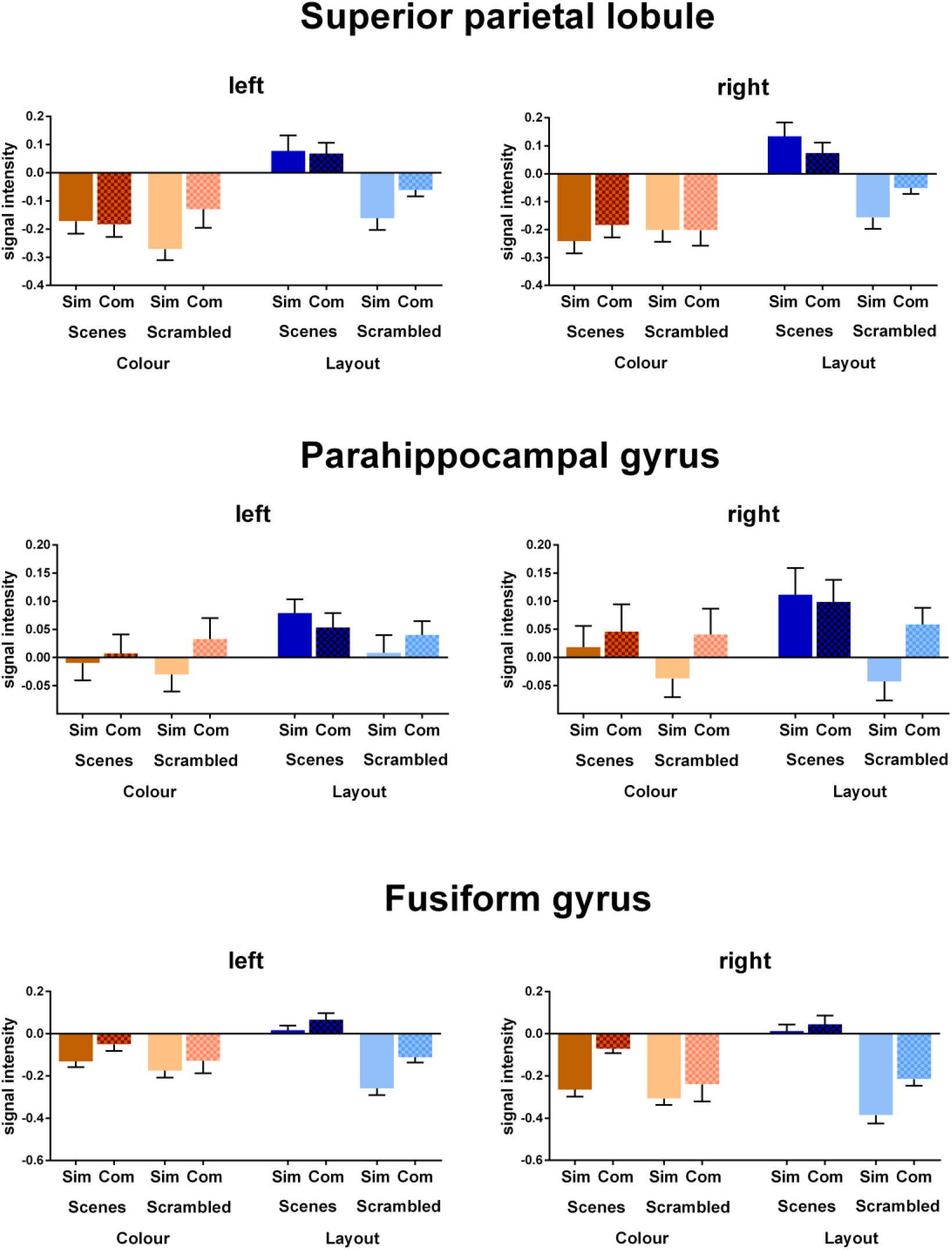
Additional signal intensities. Additional signal intensities extracted from brain regions associated with scene processing in the contrast driven PLS #1 analysis, namely the superior parietal lobule (MNI: left −20 −60 56; right 20 −60 56), parahippocampal gyrus (left −20 −15 −25; right 28 −8 −30), and fusiform gyrus (left −28 −42 −10; right 28 −42 −10). Bar graphs depict means and standard errors of the eight conditions. Sim=simple, com=complex. Of note, signal intensities are compared to an arbitrary fMRI baseline, hence negative values do not necessarily represent deactivations.

**Table S1:** Overview of the strategies used*.

| **Participant #** | **Colour conditions - simple and complex, scenes and scrambled** |
| --- | --- |
| S1 | looked at the centre of the image, compared bright areas between images |
| S2 | looked at blobs of colour, compared parts of the image but not related to the scene content |
| S3 | compared bright parts of the images |
| S4 | compared bright parts of the images |
| S5 | looked at blobs of colour, compared parts of the image but not related to the scene content |
| S6 | examined the colour along the borders of the images |
| S7 | tried to see both images at the same time |
| S8 | looked at blobs of colour, compared parts of the image but not related to the scene content |
| S9 | looked at blobs of colour, compared parts of the image but not related to the scene content |
| S10 | looked at blobs of colour, compared parts of the image but not related to the scene content |
| S11 | focused on the big picture |
| S12 | looked at blobs of colour, compared parts of the image but not related to the scene content |
| S13 | looked at blobs of colour, compared parts of the image but not related to the scene content |
| S14 | looked at blobs of colour, compared parts of the image but not related to the scene content |
| S15 | looked at blobs of colour, compared parts of the image but not related to the scene content |
| S16 | kept comparing global differences in the images |
| S17 | looked at blobs of colour, compared parts of the image but not related to the scene content |
| S18 | looked at the whole image |
| S19 | stared in the middle and so could see both images |
| S20 | looked from a distance at the whole image |
| **Participant #** | **Layout conditions - simple and complex, scenes and scrambled** |
| S1 | examined relationships between image features |
| S2 | examined relationships, focus on scenes |
| S3 | examined distance of objects from the edges |
| S4 | examined relationships between objects |
| S5 | looked at relationships, tried to see the layout of the whole picture |
| S6 | Focused on relationships between objects in scenes |
| S7 | focused on relationships between features within and between images |
| S8 | focused on relationships between features within and between images |
| S9 | focused on relationships between features within and between images |
| S10 | focused on relationships between features within and between images |
| S11 | focused on the spatial details |
| S12 | focused on relationships between features within and between images |
| S13 | focused on relationships between features within and between images |
| S14 | focused on relationships between features within and between images |
| S15 | focused on relationships between features within and between images |
| S16 | scanned around the image to see the layout |
| S17 | focused on relationships between features within and between images |
| S18 | identified most significant object, then examined their spatial relationships |
| S19 | looked horizontally and vertically, back and forth between images |
| S20 | measured and compared distance between objects |

\*Note that participants did not describe different strategies for scenes or scrambled, simple or complex images

**Table S2:** Peak coordinates of the data driven PLS LV1.

| Region | side | MNI coordinates |  |  | BSR |
| --- | --- | --- | --- | --- | --- |
|  |  | X | Y | Z |  |
| Scene construction |  |  |  |  |  |
| Middle occipital gyrus | left | -40 | -84 | 0 | 14.60 |
| Parahippocampal gyrus* | left | -30 | -30 | -20 | 9.96 |
| Precuneus* | left | -8 | -64 | 58 | 5.14 |
| Inferior parietal lobule* | left | -52 | -38 | 48 | 4.23 |
| Middle occipital gyrus | right | 36 | -82 | -6 | 14.3 |
| Fusiform gyrus* | right | 30 | -44 | -16 | 12.5 |
| Precuneus* | right | 14 | -44 | 42 | 2.92 |
| Inferior parietal lobule* | right | 50 | -44 | 50 | 2.67 |
| Inferior frontal gyrus | right | 46 | 10 | 28 | 7.84 |
| Superior parietal lobule* | right | 20 | -60 | -56 | 5.53 |
| Hippocampus (anterior)* | right | 28 | -2 | -28 | 5.23 |
| Fusiform gyrus* | right | 28 | -42 | -10 | 5.10 |
| Hippocampus (anterior)* | left | -28 | -4 | -18 | 5.02 |
| Cerebellum | left | -14 | -46 | -48 | 4.87 |
| Cerebellum | right | 20 | -42 | -46 | 4.60 |
| Insula | right | 30 | 22 | 6 | 4.06 |
| Middle cingulate gyrus | right | 8 | -2 | 32 | 4.00 |
| Middle frontal gyrus | right | 38 | 62 | 10 | 3.98 |
| Superior parietal lobule* | left | -20 | -60 | 56 | 3.67 |
| Inferior frontal gyrus | left | -40 | 36 | -18 | 3.42 |
| Ventromedial prefrontal cortex | left | -4 | 36 | -18 | 3.33 |
| Parahippocampal gyrus* | right | 28 | -8 | -30 | 3.25 |
| Fusiform gyrus* | left | -28 | -42 | -10 | 3.05 |
| All other conditions |  |  |  |  |  |
| Angular gyrus | left | -48 | -56 | 34 | 5.06 |
| Anterior cingulate gyrus | right | 4 | 32 | 0 | 4.64 |
| Angular gyrus | right | 54 | -60 | 36 | 4.55 |
| Brainstem | left | -8 | -38 | -48 | 4.27 |
| Cerebellum | right | 18 | -80 | -30 | 4.21 |
| Precuneus | left | -10 | -54 | 28 | 3.97 |
| Superior frontal gyrus | left | -12 | 36 | 48 | 3.26 |
| Middle temporal gyrus | left | -64 | -24 | -10 | 3.11 |
| Middle temporal gyrus | right | 60 | -20 | -18 | 2.88 |
| Middle cingulate gyrus |  | 0 | -22 | 40 | 2.58 |

X,Y, and Z coordinates in MNI space, BSR=Bootstrap ratio, \*=regions included in larger clusters

**Table S3:**
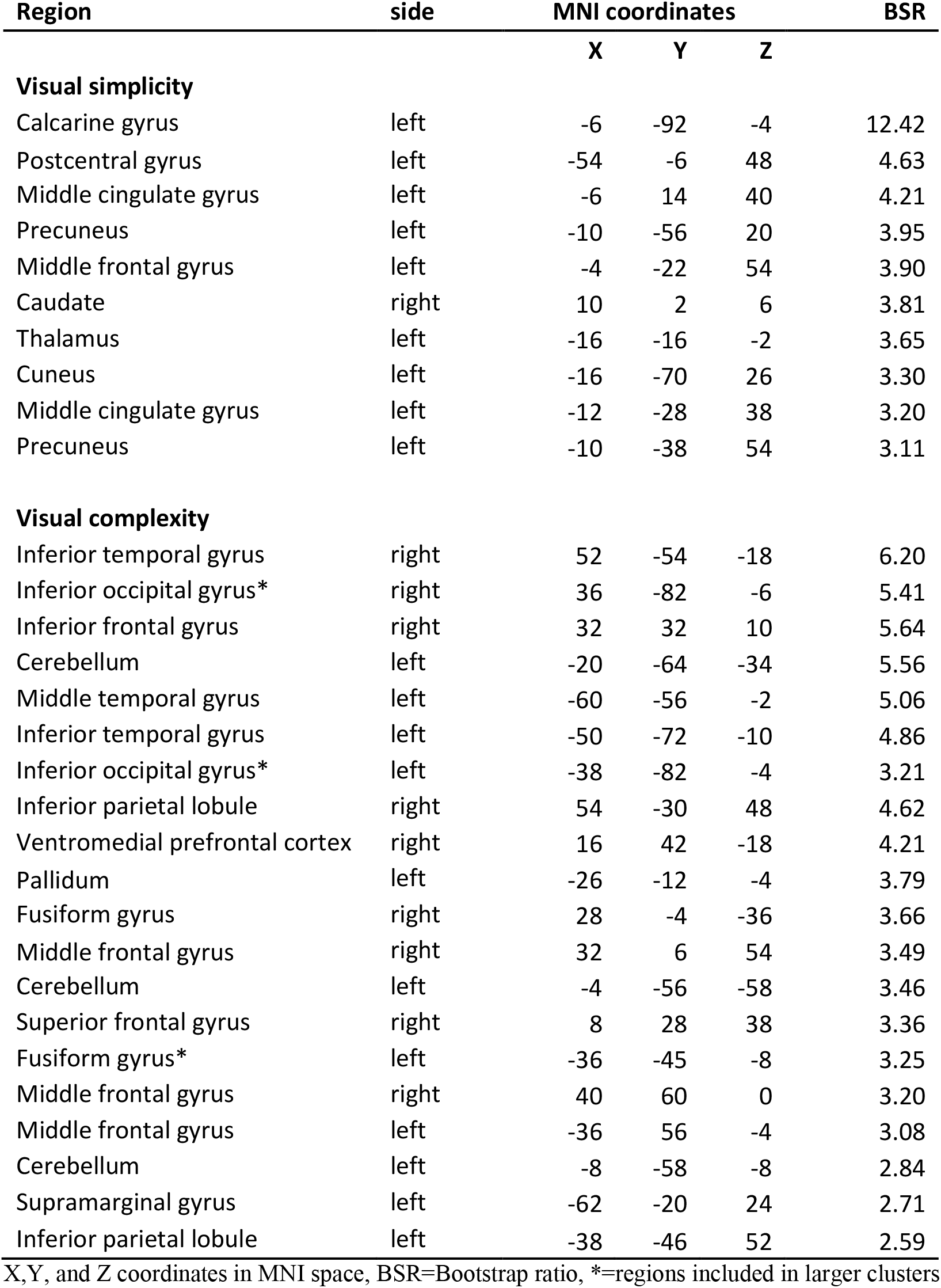
Peak coordinates of the data driven PLS LV2.

| Region | side | MNI coordinates |  |  | BSR |
| --- | --- | --- | --- | --- | --- |
|  |  | X | Y | Z |  |
| Visual simplicity |  |  |  |  |  |
| Calcarine gyrus | left | -6 | -92 | -4 | 12.42 |
| Postcentral gyrus | left | -54 | -6 | 48 | 4.63 |
| Middle cingulate gyrus | left | -6 | 14 | 40 | 4.21 |
| Precuneus | left | -10 | -56 | 20 | 3.95 |
| Middle frontal gyrus | left | -4 | -22 | 54 | 3.90 |
| Caudate | right | 10 | 2 | 6 | 3.81 |
| Thalamus | left | -16 | -16 | -2 | 3.65 |
| Cuneus | left | -16 | -70 | 26 | 3.30 |
| Middle cingulate gyrus | left | -12 | -28 | 38 | 3.20 |
| Precuneus | left | -10 | -38 | 54 | 3.11 |
| Visual complexity |  |  |  |  |  |
| Inferior temporal gyrus | right | 52 | -54 | -18 | 6.20 |
| Inferior occipital gyrus* | right | 36 | -82 | -6 | 5.41 |
| Inferior frontal gyrus | right | 32 | 32 | 10 | 5.64 |
| Cerebellum | left | -20 | -64 | -34 | 5.56 |
| Middle temporal gyrus | left | -60 | -56 | -2 | 5.06 |
| Inferior temporal gyrus | left | -50 | -72 | -10 | 4.86 |
| Inferior occipital gyrus* | left | -38 | -82 | -4 | 3.21 |
| Inferior parietal lobule | right | 54 | -30 | 48 | 4.62 |
| Ventromedial prefrontal cortex | right | 16 | 42 | -18 | 4.21 |
| Pallidum | left | -26 | -12 | -4 | 3.79 |
| Fusiform gyrus | right | 28 | -4 | -36 | 3.66 |
| Middle frontal gyrus | right | 32 | 6 | 54 | 3.49 |
| Cerebellum | left | -4 | -56 | -58 | 3.46 |
| Superior frontal gyrus | right | 8 | 28 | 38 | 3.36 |
| Fusiform gyrus* | left | -36 | -45 | -8 | 3.25 |
| Middle frontal gyrus | right | 40 | 60 | 0 | 3.20 |
| Middle frontal gyrus | left | -36 | 56 | -4 | 3.08 |
| Cerebellum | left | -8 | -58 | -8 | 2.84 |
| Supramarginal gyrus | left | -62 | -20 | 24 | 2.71 |
| Inferior parietal lobule | left | -38 | -46 | 52 | 2.59 |

X,Y, and Z coordinates in MNI space, BSR=Bootstrap ratio, \*=regions included in larger clusters

**Table S4:** Peak coordinates of the data driven PLS LV3.

| Region | side | MNI coordinates |  |  | BSR |
| --- | --- | --- | --- | --- | --- |
|  |  | X | Y | Z |  |
| Scene perception |  |  |  |  |  |
| Fusiform gyrus | right | 24 | -36 | -16 | 8.44 |
| Parahippocampal gyrus | left | -22 | -42 | -10 | 7.58 |
| Angular gyrus | left | -38 | -60 | 38 | 6.91 |
| Inferior frontal gyrus | right | 46 | 36 | 4 | 4.82 |
| Posterior cingulate gyrus | left | -2 | -36 | 40 | 4.65 |
| Hippocampus (posterior)* | left | -36 | -28 | -14 | 4.55 |
| Fusiform gyrus* | left | -36 | 54 | -10 | 4.27 |
| Supramarginal gyrus | left | -58 | -20 | 20 | 4.06 |
| Postcentral gyrus | left | -52 | -18 | 54 | 3.65 |
| Inferior occipital gyrus* | right | 38 | -76 | -8 | 3.52 |
| Inferior temporal gyrus | right | 34 | 0 | -38 | 3.42 |
| Inferior frontal gyrus | right | 58 | 12 | 10 | 3.20 |
| Inferior occipital gyrus* | left | -34 | -76 | -10 | 2.83 |
| Parahippocampal gyrus* | right | 28 | -26 | -20 | 2.57 |
| Hippocampus (posterior)* | right | 32 | -32 | -4 | 2.47 |
| Scrambled construction |  |  |  |  |  |
| Dorsomedial prefrontal cortex | right | 12 | 38 | 22 | 6.52 |
| Superior parietal lobule | right | 16 | -70 | 58 | 5.30 |
| Cerebellum | right | 36 | -68 | -46 | 5.06 |
| Middle cingulate cortex | left | -8 | 14 | 44 | 4.80 |
| Inferior frontal gyrus | right | 54 | 10 | 24 | 4.79 |
| Brainstem | left | -4 | -18 | -12 | 4.46 |
| Middle temporal gyrus | right | 50 | -24 | -16 | 4.11 |
| Middle frontal gyrus | left | -36 | -2 | 44 | 4.11 |
| Superior parietal lobule | left | -18 | -56 | 60 | 3.96 |
| Cerebellum | left | -14 | -46 | -58 | 3.73 |
| Cerebellum | right | 6 | -58 | -52 | 3.72 |
| Inferior parietal lobule | right | 40 | -38 | 52 | 3.35 |
| Caudate | right | 10 | 6 | 8 | 3.12 |
| Postcentral gyrus | right | 66 | 0 | 30 | 2.60 |

X,Y, and Z coordinates in MNI space, BSR=Bootstrap ratio, \*=regions included in larger clusters

**Table S5:** Peak coordinates of the contrast driven PLS - scenes versus scrambled images.

| Region | side | MNI coordinates |  |  | BSR |
| --- | --- | --- | --- | --- | --- |
|  |  | X | Y | Z |  |
| Naturalistic scenes |  |  |  |  |  |
| Middle occipital gyrus | left | -38 | -78 | 16 | 19.02 |
| Fusiform gyrus* | left | -30 | -44 | -16 | 15.13 |
| Lingual gyrus* | right | 26 | -64 | -12 | 12.29 |
| Lingual gyrus* | left | -24 | -62 | -10 | 11.31 |
| Parahippocampal gyrus* | right | 24 | -30 | -18 | 9.69 |
| Superior frontal gyrus | left | -24 | 12 | 58 | 6.80 |
| Fusiform gyrus | right | 32 | -2 | -36 | 6.03 |
| Precentral gyrus | right | 40 | 2 | 56 | 5.44 |
| Parahippocampal gyrus* | left | -36 | -18 | -22 | 4.84 |
| Middle frontal gyrus | right | 52 | 48 | -6 | 4.57 |
| Inferior parietal lobule* | right | 28 | -62 | 42 | 4.27 |
| Hippocampus (anterior)* | right | 18 | -8 | -22 | 4.27 |
| Hippocampus (posterior)* | right | 23 | -36 | 2 | 3.28 |
| Brainstem | left | -8 | -34 | -36 | 3.64 |
| Ventromedial prefrontal cortex | right | 2 | 50 | -22 | 3.63 |
| Middle cingulate gyrus | right | 8 | -2 | 32 | 3.58 |
| Middle frontal gyrus | left | -46 | 50 | -4 | 3.46 |
| Globus Pallidus | left | -14 | -2 | 4 | 3.42 |
| Cerebellum | left | -30 | -68 | -50 | 3.19 |
| Rolandic operculum | left | -50 | -22 | 18 | 3.01 |
| Superior temporal pole | right | 50 | 20 | -16 | 2.89 |
| Hippocampus (anterior)* | left | -24 | -10 | -20 | 2.71 |
| Hippocampus (posterior)* | left | -18 | -31 | -2 | 2.67 |
| Inferior parietal lobule* | left | -50 | -44 | 50 | 2.57 |
| Middle frontal gyrus | right | 26 | 36 | -14 | 2.52 |
| Scrambled images |  |  |  |  |  |
| Precuneus | left | -14 | -54 | 30 | 4.31 |
| Anterior cingulate gyrus | right | 2 | 30 | 0 | 3.82 |
| Cerebellum | right | 2 | -60 | -48 | 3.71 |
| Calcarine gyrus | left | -12 | -92 | 4 | 3.49 |
| Middle frontal gyrus | right | 26 | 48 | 36 | 3.36 |
| Middle temporal gyrus | right | 52 | -28 | -10 | 3.34 |
| Middle cingulate gyrus | left | -2 | -24 | 30 | 2.99 |
| Superior frontal gyrus | right | 12 | 42 | 22 | 2.91 |
| Superior frontal gyrus | left | -6 | 60 | 8 | 2.32 |

X,Y, and Z coordinates in MNI space, BSR=Bootstrap ratio, \*=regions included in larger clusters

**Table S6:** Peak coordinates of the contrast driven PLS - scene construction versus scene perception.

| Region | side | MNI coordinates |  |  | BSR |
| --- | --- | --- | --- | --- | --- |
|  |  | X | Y | Z |  |
| Scene construction |  |  |  |  |  |
| Superior parietal lobule | right | 16 | -74 | 58 | 8.16 |
| Precentral gyrus | right | 32 | 0 | 54 | 7.97 |
| Cerebellum | right | 36 | -70 | -46 | 6.33 |
| Fusiform gyrus* | right | 34 | -38 | -16 | 5.86 |
| Fusiform gyrus | left | -42 | -60 | -14 | 5.73 |
| Hippocampus (anterior) | left | -32 | -2 | -22 | 5.08 |
| Middle occipital gyrus* | right | 34 | -80 | 26 | 4.97 |
| Superior frontal gyrus | right | 6 | 62 | 36 | 4.89 |
| Lingual gyrus* | left | -24 | -48 | -10 | 4.61 |
| Superior occipital gyrus* | left | -28 | -82 | 34 | 4.34 |
| Cerebellum | left | -30 | -46 | -38 | 4.16 |
| Ventromedial prefrontal cortex | left | -8 | 44 | -22 | 4.08 |
| Hippocampus (anterior) | right | 24 | -8 | -24 | 3.81 |
| Parahippocampal gyrus* | right | 28 | -22 | -24 | 3.75 |
| Middle frontal gyrus | left | -38 | 36 | -16 | 3.54 |
| Parahippocampal gyrus* | left | -22 | -16 | -22 | 3.21 |
| Inferior frontal gyrus | right | 30 | 46 | -18 | 3.04 |
| Scene perception |  |  |  |  |  |
| Angular gyrus | left | -48 | -60 | 42 | 6.60 |
| Anterior cingulate gyrus | right | 2 | 32 | 2 | 4.45 |
| Superior frontal gyrus | right | 18 | 30 | 38 | 3.91 |
| Angular gyrus | right | 54 | -54 | 50 | 3.89 |
| Precuneus | right | 14 | -50 | 34 | 3.72 |
| Hippocampus (posterior) | left | -28 | -36 | 4 | 3.67 |
| Hippocampus (body) | left | -34 | -18 | -14 | 3.57 |
| Insula | left | -48 | 12 | 6 | 3.57 |
| Superior frontal gyrus | left | -12 | 28 | 58 | 3.33 |
| Middle temporal gyrus | left | -60 | -22 | -6 | 2.77 |
| Middle frontal gyrus | left | -32 | 22 | 44 | 2.75 |

X,Y, and Z coordinates in MNI space, BSR=Bootstrap ratio, \*= in the text referred to as ventromedial prefrontal cortex (vmPFC).

